# Selective whole-genome sequencing of *Plasmodium* parasites directly from blood samples by Nanopore adaptive sampling

**DOI:** 10.1101/2022.11.29.518068

**Authors:** Katlijn De Meulenaere, Wim L. Cuypers, Anna Rosanas-Urgell, Kris Laukens, Bart Cuypers

## Abstract

**Background:** Whole-genome sequencing (WGS) is becoming an increasingly popular tool to study the population genetics and drug resistance of *Plasmodium* spp. However, the predominance of human DNA in a malaria patient blood sample requires time-consuming lab procedures to filter out human DNA or enrich *Plasmodium* DNA. Here, we investigated the potential of adaptive sampling to enrich for *Plasmodium* DNA while sequencing unenriched patient blood samples on a minION device.

**Results:** To compare adaptive sampling versus regular sequencing, a dilution series consisting of 0% up to 100% *P. falciparum* DNA in human DNA was sequenced. Half of the flowcell channels were run in adaptive sampling mode, enriching for the *P. falciparum* reference genome, resulting in a 3.2 fold enrichment of *P. falciparum* bases on average. Samples with a lower concentration of parasite DNA had a higher enrichment potential. We confirmed these findings by sequencing two *P. falciparum* patient blood samples with common levels of parasitaemia (0.1% and 0.2%). The estimated enrichment was 3.9 and 5.8, which was sufficient to cover at least 97% of the *P. falciparum* reference genome at a median depth of 20 (highest parasitaemia) or 5 (lowest parasitaemia). A comparison of 38 drug resistance variants (WHO) obtained via adaptive sequencing or Sanger sequencing showed a high concordance between the two methods, suggesting that the obtained sequencing data is of sufficient quality to address common clinical research questions for patients with parasitaemias of 0.1% and higher.

**Conclusions:** Our results demonstrate that adaptive Nanopore sequencing has the potential to replace more time-consuming *Plasmodium*-enrichment protocols and sequence directly from patient blood, given further improvements in cost-efficiency.

## Introduction

The large majority of human malaria cases are caused by the *Plasmodium falciparum* parasite. In recent years, the downward trend in the number of cases per year is stagnating (WHO world malaria report 2021). A better understanding of the population dynamics of *P. falciparum* is therefore required. In addition, increasing reports of drug resistance and deletions in the rapid diagnostic test (RDT) target genes *pfhrp2/3*, warrant genomic surveillance of *Plasmodium*^1,2^. The most comprehensive method to simultaneously study population dynamics and characterize resistance determinants is whole-genome sequencing (WGS).

It remains challenging to obtain sufficient amounts of parasite DNA straight from patient blood samples to fulfill WGS requirements, and multi-step wet-lab procedures are required^3–5^. A blood sample of a patient infected with *Plasmodium* consists of infected red blood cells containing malaria DNA, non-infected red blood cells (RBCs) that contain no DNA (no nucleus), and white blood cells (WBCs) that contain human DNA. A low fraction of the red blood cells are infected, typically <2.5% for patients with uncomplicated *P. falciparum* malaria^6^. Therefore, the majority of the obtained DNA will be human. To avoid expensive futile sequencing of human DNA, leukocyte depletion and selective whole genome amplification (sWGA) are commonly used^3,4^. Leukocyte depletion techniques should be applied to freshly collected blood samples, which limits the applicability of this technique in low-resource or remote settings where laboratory facilities are not often close to the site of collection^7^. sWGA can be performed on frozen blood samples or dried blood spots, but is more expensive, time-consuming, and potentially introduces amplification bias^3,4^.

Ideally, WGS is carried out on fresh samples and in endemic settings where patients would benefit directly from clinical genomics. Sequencing devices of Oxford Nanopore Technologies (ONT) have enabled mobile and real-time sequencing ^8,9^. During Nanopore sequencing, DNA is translocated via a protein pore through which an ionic current flows. The DNA molecule disturbs this current, generating an electrical signature that corresponds to a set of nucleotides and their modifications^10^. Because the voltage across the pore can be reversed, molecules can also be ejected. Signals originating from the DNA molecules that are being sequenced can be classified in real time and used to decide whether a molecule should be ejected or not. This procedure is referred to as ‘adaptive sampling’ and allows the enrichment of a target of interest while sequencing^11–14^.

In this paper, we assessed the utility of adaptive sampling for the enrichment of *Plasmodium* DNA in blood samples from human *P. falciparum* patients, without a prior leukocyte depletion or parasite enrichment step. Two experiments were performed (**Figure 1**). In the first experiment, we compared adaptive sampling to regular sequencing for a dilution series of human DNA mixed with *Plasmodium* DNA. In the second experiment, we sequenced two patient samples using the adaptive sampling feature and assessed the quality of the obtained data in light of clinical applications. Our results demonstrate that adaptive sampling enables a sufficiently deep sequencing of patient samples with commonly occurring parasitaemias (0.1-0.2%).

**Figure 1.**
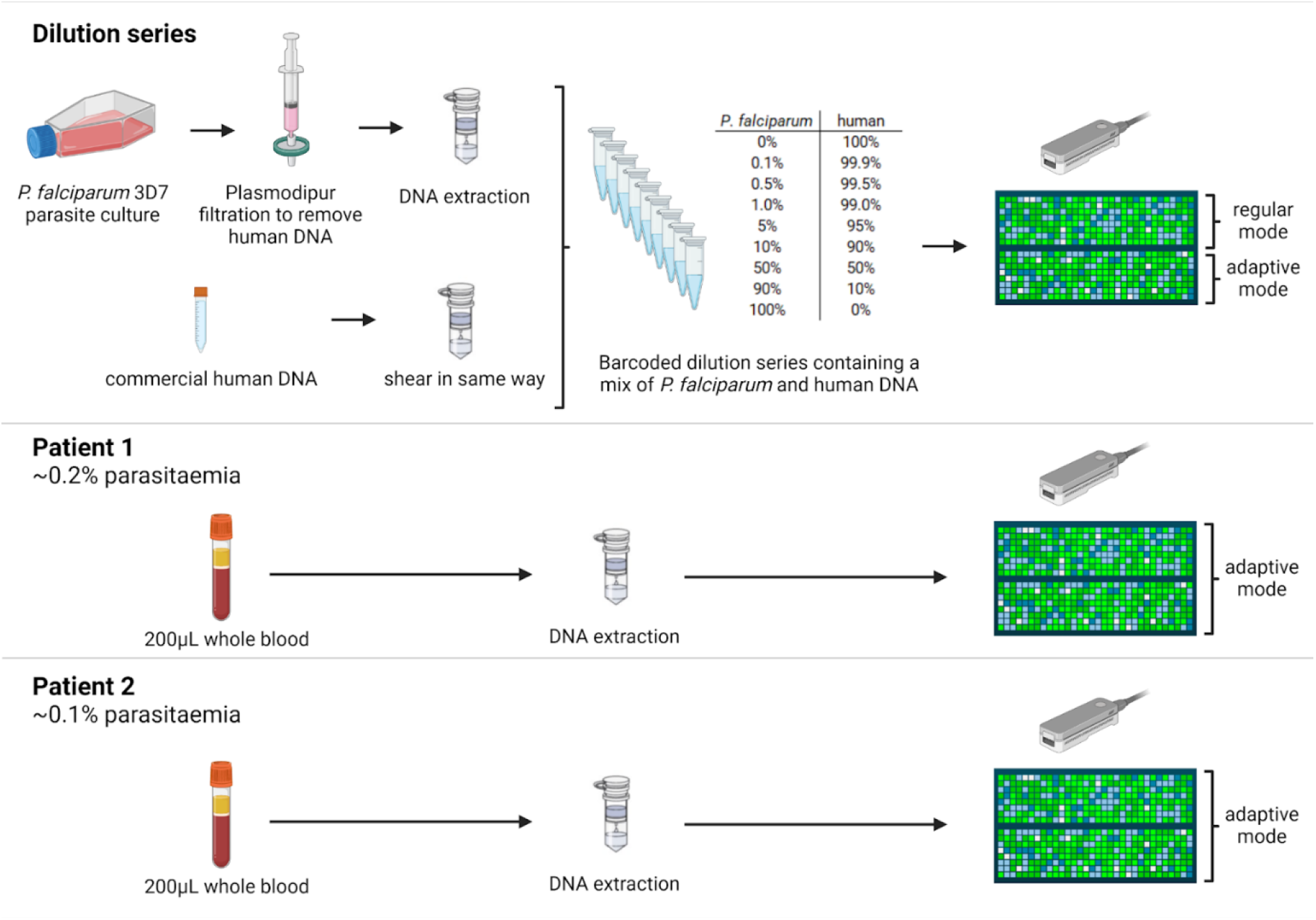
Overview of the experimental setup. First, a dilution series of human DNA mixed with *P. falciparum* DNA was sequenced with half of the ONT flowcell operating in adaptive mode and the other half in regular mode. Next, two *P. falciparum* patient samples were sequenced in adaptive mode, without prior leukocyte depletion or parasite enrichment.

## Materials and Methods

### Samples and DNA extraction

A dilution series of simulated patient samples containing 0%, 0.1%, 0.5%, 1%, 5%, 10%, 50%, 90% and 100% *P. falciparum* DNA was made by mixing *P. falciparum* 3D7 DNA with commercially available human DNA (Promega, G304A) to a total amount of 400ng DNA. *P. falciparum* strain 3D7 was *in vitro* cultured to 5-8% parasitaemia, after which the parasites were harvested and remaining human white blood cells were depleted with Plasmodipur filtration (EuroProxima) as described by Auburn et al. (2011)^3^. DNA was extracted from 400μL RBCs using the QIAamp DNA Mini kit (Qiagen). The commercial human DNA was processed using the same QIAamp DNA Mini columns to obtain similar DNA fragment sizes for both the human and *P. falciparum* DNA. Fragment sizes were assessed with the Fragment Analyser (Agilent, kit DNF-464-33 -HS Large Fragment 50Kb) (**Supplementary Figure 1**).

Clinical samples originated from two patients who presented themselves at the Institute of Tropical Medicine Antwerp (ITM) travel clinic with a *P. falciparum* infection. DNA was extracted from 200μL whole blood (stored in EDTA) using the QIAamp DNA Mini kit (Qiagen). Parasite density (parasites/μL) was determined by varATS qPCR, based on a protocol adapted from Hofmann et al. (2015)^15^. Sample 1 contained 11687 parasites/μL, which corresponds to a parasitaemia of approximately 0.23% (infected RBCs/RBCs). Sample 2 contained 4508 parasites/μL, which corresponds to approximately 0.09% parasitaemia.

### Library preparation and sequencing

The Human and *Plasmodium* DNA dilution series consisted of nine mixed samples that were barcoded and prepared for sequencing using the Native Barcoding Kit 24 (Q20+) (SQK-NBD112.24) following the manufacturer’s protocol (version NBE_9134_v112_revE_01Dec2021). The resulting barcoded library was loaded on one R10.4 flowcell, and sequencing was performed on a MinION sequencer with MinKnow v23.03.6 (ONT). Channels 1 - 256 were operating in adaptive sampling mode, with the 3D7 reference genome (PlasmoDB v56) listed as the target genome for enrichment. Channels 257 - 512 were operating in regular sequencing mode. After 72 hours, 4.26 gigabase (Gb) of data was generated and the run was stopped.

Two additional libraries were prepared from clinical samples 1 and 2 using the Ligation Sequencing Kit V14 (SQK-LSK114 - Early Access product at the time of writing). The manufacturer’s protocol was followed. The same two patient samples were also sequenced before with the SQK-LSK112 kit, but because kit SQK-LSK114 clearly delivered higher output and therefore better *P. falciparum* sequencing depths (comparison in **Supplementary Table 1**), only SQK-LSK114 data will be shown and discussed.

For patient samples 1 and 2, 586ng and 498.2ng of DNA was used for library preparation respectively, and the library was split into two parts. Sequencing was performed on a MinION sequencer (R10.4.1 flowcell) coupled to MinKnow v22.08.9 (ONT). The sequencing speed was set at 400bps. The flowcell was washed with the Flow Cell Wash Kit (EXP-WSH004) after 33 hours for the first sample and 37 hours for the second sample, after which the second part of the library was loaded. The run was stopped after 80 and 70 hours resulting in 9.41 and 9.74 gigabase (Gb) output, respectively.

### Basecalling and species identification

Basecalling was performed offline using Guppy GPU v6.0.7 for the dilution series, Guppy GPU v6.3.8 for patient sample 1, and Guppy GPU v6.2.11 for patient sample 2. All data was basecalled using the super-accurate model. Quality metrics were assessed using pycoQC v2.5.2^16^ and Nanoplot v1.28.1^17^. For the dilution series experiment, reads were separated according to sequencing mode (adaptive sampling versus regular sequencing) using seqtk v1.3^18^.

Reads were competitively mapped using minimap2 v2.17-r941 to a reference fasta file consisting of the human (GRCh38) and *P. falciparum* 3D7 (PlasmoDB v58) reference genomes, to minimize false-positive mapping. Unmapped (bitwise flag 4) and alternatively mapping (bitwise flag 256) reads were removed with samtools v1.10 from coordinate sorted bam files, after which *P. falciparum* and human reads could be split based on their genome-specific coordinates. The number of mapped bases was extracted from the samtools stats report (“bases mapped (cigar)”), the number of mapped reads from the samtools flagstat report. Depth per position was obtained with the samtools depth command, and further analyzed in R^19^. Coverage was calculated as the fraction of the reference genome that was covered by at least one uniquely mapping read. Depth was calculated as the average number of reads that cover a position in the reference genome.

Taxonomic read classification was performed using kraken2^20^ combined with the PlusPF-16 database (retrieved from https://benlangmead.github.io/aws-indexes/k2 on 10/05/2022) to assess the species composition of the samples and check for potential contaminants.

Reads of the dilution series’ positive control sample (100% human DNA) that mapped to the *P. falciparum* reference genome were subjected to an additional analysis using blastn v2.13.0+^21,22^ to confirm whether these reads originated from *P. falciparum* (Plasmodipur filtration is not 100% efficient) or showed homology to the human reference genome. Reads were first converted to the fasta format using seqtk^18^ and compared to a custom database consisting of *Homo sapiens* genome assembly GRCh38.p14 and *P. falciparum* GCA_000002765.3 (retrieved on 03/08/2022). For reads with hits to the human reference genome, a maximum of five hits with the highest bitscore were extracted and further inspected.

### Qualitative comparison of sequencing output between both *P. falciparum* patient samples

To estimate the extent to which adaptive sampling enriched *P. falciparum* DNA in the clinical samples, we compared the fraction of sequenced *P. falciparum* bases to the sequenced fraction of *P. falciparum* reads. The latter is an approximation of the unknown fraction of *P. falciparum* DNA that is present in the patient samples. To take into account that adaptive sampling decreases the total yield compared to regular mode sequencing^11^, this number is divided by 1.6 (yield correction factor determined via the dilution series experiment) to get the *P. falciparum* enrichment in comparison to a run in regular mode.

Variants of the clinical samples were called with Medaka v1.6.1 (medaka_haploid_variant, model r1041_e82_400bps_sup_variant_g615) (https://github.com/nanoporetech/medaka). To validate the called variants, a set of genes with known drug resistance markers (*pfdhfr, pfmdr1, pfcrt, pfK13, pfcytb*) was PCR amplified and Sanger sequenced (Genewiz, Germany) according to existing methods^23–26^. Multiplicity of infection (MOI), *i*.*e*. a polyclonal infection, was determined by MSP1 and MSP2 genotyping^27^.

*De novo* genome assembly of *P. falciparum* reads originating from patient 1 and 2 was carried out using Flye v2.9^28^ with the option --nano-hq and the --genome-size parameter set to 23m. One round of polishing based on the *P. falciparum* mapped reads was carried out with Medaka v1.6.1 (medaka_consensus, model r1041_e82_400bps_sup_variant_g615) (https://github.com/nanoporetech/medaka). Scaffolding of contigs against the *P. falciparum* 3D7 reference (PlasmoDB v58) was performed using RagTag v2.1.0, without prior correction to maintain true biological structural variation. The resulting scaffolds were annotated with Companion v1.0.2^29^.

## Results

### Adaptive sampling enriches *P. falciparum* DNA 3.2-fold

To test whether and to which degree adaptive sampling can enrich low-abundant *P. falciparum* DNA in a human blood sample, we made a series of nine *P. falciparum* DNA samples diluted in commercial human DNA at different percentages, ranging from 0% *P. falciparum* and 100% human to 100% *P. falciparum* and 0% human (**Figure 1**). Enrichment by adaptive sampling depends on the fragment size of the input DNA^11^, which was determined using a Fragment Analyzer for the 0% and 100% *P. falciparum* samples. The average fragment size for the human sample was circa 39kb, while the *P. falciparum* DNA peaked at around 46kb (fragment size distributions in **Supplementary Figure 1**). Samples were pooled and sequenced on one flowcell with half of the channels operating in adaptive sampling mode (enriching for the *P. falciparum* reference genome), and half of the channels operating in regular sequencing mode (**Figure 1**).

Adaptive sampling enriched the number of bases mapping to the *P. falciparum* reference genome 3.2 fold on average (range of 0.7 to 5.1 fold) for the different concentrations (**Figure 2A-B**). Lower overall yield was previously observed during sequencing runs in adaptive sampling mode^11,13^. Hence, we calculated the enrichment by dividing the fraction of *P. falciparum* bases sequenced in adaptive mode by the fraction of *P. falciparum* bases in regular mode, while normalizing for the lower output in adaptive mode compared to regular mode (1.6x lower for this run).

**Figure 2.**
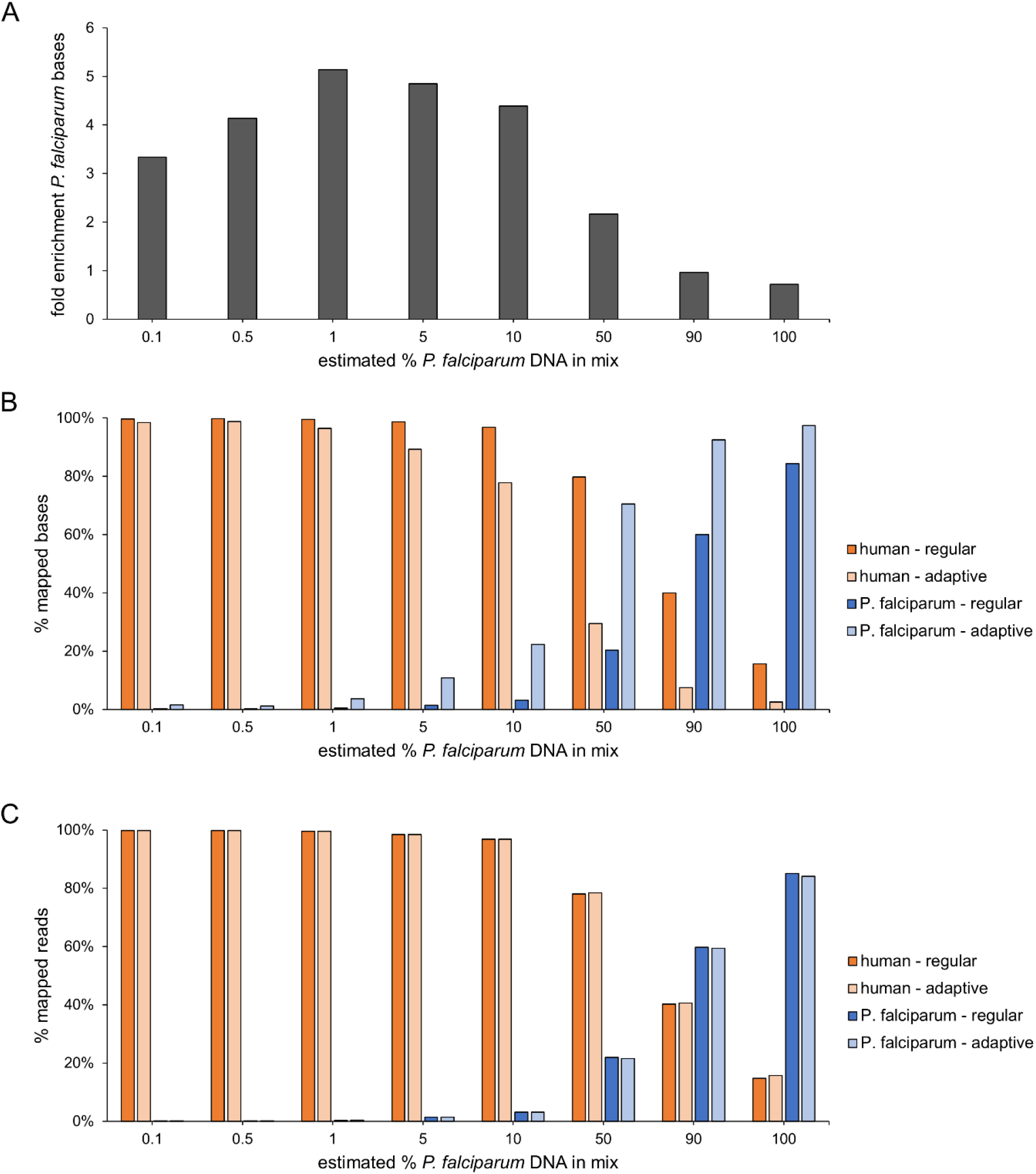
Comparison of human and *P. falciparum* reads, bases and enrichment for regular and adaptive sampling. **A**. Enrichment for *P. falciparum* DNA in adaptive sampling mode. Bars indicate the enrichment for bases mapping to the *P. falciparum* reference genome, taking into account the lower yield in adaptive mode. **B**. Adaptive sampling increased the number of mapped on-target (*P. falciparum*) bases and decreased the number of off-target (human) bases. The sum of human and *P. falciparum* % of reads equals 100% per sequencing mode. **C**. Percentages of reads per species in adaptive sampling mode versus regular sequencing mode. The sum of human and *P. falciparum* % of reads equals 100% per sequencing mode.

Further inspection of the read lengths and read fractions per species provided mechanistic insight in the adaptive sampling procedure. The percentage of reads per species remained unaltered (**Figure 2C**), human reads were ejected after sequencing approximately 400 bases, which allowed at least three times more reads to be sequenced (**Supplementary Figure 2, Supplementary Table 2**). In adaptive sampling mode, the median read length was 391bp for human reads and 1701bp for *P. falciparum* reads (**Supplementary Figure 2**).

Samples with low percentages *P. falciparum* DNA (0.1%, 0.5%, 1, 5% and 10%) showed the highest enrichment (**Figure 2A**). In comparison, the fold enrichment was half or less in samples where *P. falciparum* DNA was more abundant (50%, 90% and 100%) (**Figure 2A**). Due to the higher sequencing efficiency in regular mode, the sample with the highest amount of *P. falciparum* DNA had a fold enrichment of 0.7, showing that the output was higher in the regular sequencing mode. Adaptive sampling substantially increases the amount of sequenced *P. falciparum* bases when sequencing low parasitaemia samples, but loses efficiency around 50% *P. falciparum* DNA, and highly enriched samples with >90% of *P. falciparum* DNA can be more efficiently sequenced in the regular sequencing mode.

A positive control sample consisting exclusively of commercial human DNA and 0% *P. falciparum* DNA was added to estimate the degree of barcode contamination. Of all reads in the positive control sample, 0.17% mapped to the *P. falciparum* reference genome, corresponding to 0.61% of the total amount of bases sequenced for this sample. Homology between these reads and the human reference genome was rare as determined using blastn. Out of 33 and 114 reads that were identified as *P. falciparum* in regular- and adaptive sampling mode respectively, only 5 had secondary hits to the human reference genome (**Supplementary Table 3**). This indicates that the presence of *P. falciparum* reads in the human sample likely resulted from barcode contamination. Conversely, our experiment did not allow an assessment of potential barcode contamination with human reads in the 100%

*P. falciparum* sample because the Plasmodipur filter removal of human white blood cells is not 100% efficient. Indeed, 15.7% of the reads in the sample consisting of 100% purified *P. falciparum* DNA were identified as human.

### Direct adaptive sequencing of patient blood samples

Patient samples 1 and 2 were each sequenced on one MinION flowcell (**Figure 1**). Enrichment for the *P. falciparum* reference genome via adaptive sampling was enabled. Taxonomic read classification revealed that more than 98.89% and 99.59% of sequencing reads were human. Most of the remaining reads originated from *P. falciparum* and *Toxoplasma gondii* which is highly prevalent in humans (**Supplementary Table 4**). Opposed to patient sample 1, patient sample 2 contained less *P. falciparum* reads than *T. gondii* reads (**Supplementary Table 4**).

Competitive read mapping and subsequent analysis suggest that for patient samples with commonly occurring parasitaemias (∼0.1%-0.2%), sufficient coverage and sequencing depth can be obtained for the entire *P. falciparum* 3D7 reference genome with adaptive sampling. Reads from patient sample 1 with the highest parasitaemia of 0.23% covered 99.8% of the *P. falciparum* 3D7 reference genome with a median depth of 20. The majority (93.4%) of the sequenced bases were human, and 6.7% of the bases and 1.1% of the reads were *P. falciparum*. For patient sample 2, 1.7% of the sequenced bases and 0.2% of the reads were identified as *P. falciparum*. This resulted in a median depth of 5, and a *P. falciparum* genome coverage of 97.4%. These results are in line with patient 2’s lower parasitaemia of 0.09%. Sequencing depth was uniform across all chromosomes. The mitochondrial genome however stands out with a median depth of 522 and 125 for patient 1 and 2 respectively, due to the presence of multiple plasmid copies (**Supplementary Tables 5 and 6**).

In line with our observation that more diluted target DNA becomes more enriched by adaptive sampling, we found that adaptive sampling successfully enriched the amount of sequenced parasite DNA 3.9 fold for patient sample 1, and 5.8 fold for patient sample 2. This estimate was corrected for the lower yield of a flowcell operating in adaptive sampling mode which we observed for the dilution series experiment. As a result, this estimate reflects the enrichment compared to a (higher total output) run in regular sequencing mode.

Several downstream analyses were conducted to assess whether the data was of sufficient quality to address common clinical and/or research questions. *De novo* genome assembly of *P. falciparum* reads showed that fragmented assemblies could be obtained. For patient sample 1, the assembly size was slightly larger than the current 3D7 genome size (105%) and 138 contigs were obtained, while the assembly size of patient sample 2 was 95% of the reference genome size and 198 contigs were obtained. The higher sequencing depth of patient 1 resulted in a less fragmented assembly, where chromosomes consisted of only 1 up to 9 contigs, while in patient 2, 6 to 26 contigs were assigned to each chromosome (**Supplementary Table 7**). The apicoplast and mitochondrial plasmid in both cases consisted of one contig that entirely covered the reference sequences of the apicoplast and mitochondrial plasmid of the *P. falciparum* 3D7 reference genome (**Supplementary Table 7**). Although the *de novo* assemblies were fragmented, many coding genes could be detected, 5323 (100.1%) and 4733 (89.0%) for patient 1 and 2 respectively.

To assess whether direct sequencing of patient samples with adaptive sampling yields data of sufficient quality for SNP calling (*e*.*g*. to screen for SNPs implicated in drug resistance or inferring relatedness to other *Plasmodium* strains), we screened the sequencing data for 57 drug resistance-associated markers^31^ and compared the results to Sanger sequencing data of 38 of these markers (**Supplementary Table 8**). Overall, there was a high concordance of 97% for patient 1 and 100% for patient 2 between the results obtained through Nanopore sequencing and Sanger sequencing (**Table 1**). For patient 1, we found 5 sulfadoxine-resistance associated SNPs out of the 57 markers that were all sequenced confidently (depth >10). Because patient 1 has a polyclonal infection (MOI=4), minority clones carrying drug resistant alleles were identified for multiple loci (**Supplementary Table 8**). This could have caused the single mismatch between the Nanopore and Sanger sequencing result at the *pfcrt* K76T locus. For patient 2 (MOI=1), 4 drug resistance SNPs were found **(Table 1**), and although sequencing depth is lower **(Supplementary Table 8**), Sanger sequencing confirms that their variants were correctly called. Because long Nanopore reads are well suited to resolve structural variants, 2 rapid diagnostic test (RDT) target genes (*pfhrp2, pfhrp3*) known to show large deletions were investigated^2^. *Pfhrp3* showed consistent deletions with clear boundaries in the 3th exon for both patients.

**Table 1.** Concordance and discordance between Nanopore (adaptive mode) and Sanger sequencing results of 57 WHO drug resistance markers^31^ in two *P. falciparum* patient samples. MOI=multiplicity of infection; P=parasitaemia.

| PATIENT 1 (MOI=4, ~0.2%P) |  |  |  |
| --- | --- | --- | --- |
|  | Nanopore result | Sanger result | drug-resistance markers |
| Concordant | Sensitive | Sensitive | 33 |
|  | Resistant | Resistant | 4 |
| Discordant | Sensitive | Resistant | 1 |
|  | Resistant | Sensitive | 0 |
| Missing data | Sensitive | NA | 18 |
|  | Resistant | NA | 1 |

| PATIENT 2 (MOI=1, ~0.1%P) |  |  |  |
| --- | --- | --- | --- |
|  | Nanopore result | Sanger result | drug-resistance markers |
| Concordant | Sensitive | Sensitive | 35 |
|  | Resistant | Resistant | 3 |
| Discordant | Sensitive | Resistant | 0 |
|  | Resistant | Sensitive | 0 |
| Missing data | Sensitive | NA | 18 |
|  | Resistant | NA | 1 |

### Sequencing in adaptive mode is faster and more flexible compared to existing but cheaper methods

We compared the cost of our method to the current state-of-the-art for *Plasmodium* WGS that relies on additional enrichment steps. The cost of sequencing one unenriched blood sample on a MinION flowcell was approximately 1000 euro when we conducted this study. In comparison, common methods used to sequence a *P. falciparum* genome in its entirety are much more cost-efficient (*e*.*g*. 100 euro), based on our *in-house* costs for sWGA and leukocyte depletion, and the publication of Melnikov et al. (2011) on hybrid selection^5^ (**Figure 3**).

**Figure 3.**
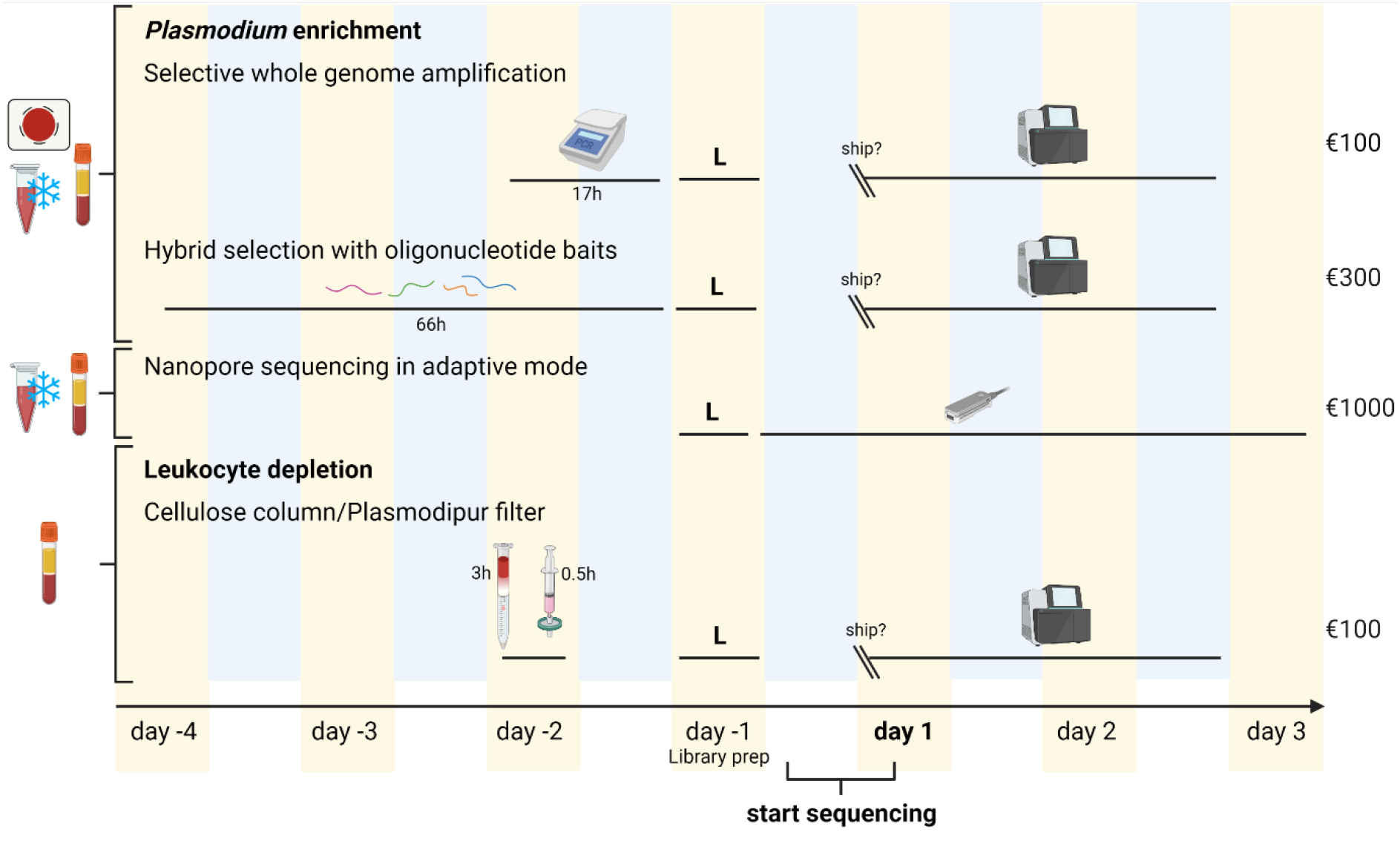
Overview of commonly used methods to increase the amount of *Plasmodium* reads vs human reads, by enrichment of *Plasmodium* DNA or leukocyte depletion, and comparison of duration and cost with nanopore adaptive sequencing. For leukocyte depletion methods, only a fresh whole blood sample can be used, while other methods can also be carried out on frozen blood or a dried blood spot (indicated on the left). On the right, an estimate of the full cost per sample (method-specific preparations, library preparation, sequencing; cost at the moment of writing) is given. ‘L’ stands for library preparation; yellow and blue background colors indicate day and night, respectively.

In contrast to leukocyte depletion methods, the adaptive sequencing workflow can also start from frozen blood samples, which improves field applicability. The same applies for the other parasite enrichment methods shown in **figure 3** (sWGA, hybrid selection), but those require a more lengthy sample preparation in comparison to the nanopore procedure, which can be done straight from whole blood.

The time required to process a sample and obtain *P. falciparum* sequence data is up to four days using Nanopore adaptive sampling, which is shorter compared to the commonly used methods for *P. falciparum* DNA enrichment and sequencing (**Figure 3**). Starting from extracted DNA, a library can be prepared and loaded on a MinION flowcell within 2 hours. Sequencing takes up to three days to reach the highest depth, but since the bulk of the data is generated during the first day, variants could be called earlier, and rapid identification of the infecting *Plasmodium* species could already be achieved within a couple of hours. Nanopore basecalling can be done during the run, but is preferably carried out afterwards^32^, which takes up to 1-1.5 additional days.

## Discussion

We investigated the feasibility of using adaptive sampling to enrich for *Plasmodium* DNA while sequencing unenriched patient blood samples on a minION device. We showed that on average a 3.2 fold enrichment can be achieved for samples with different concentrations of *P. falciparum* DNA and that samples with a lower concentration of parasite DNA (<50%) have a higher enrichment potential. For highly purified parasite samples with over 90% *P. falciparum* DNA, regular sequencing yields the highest output of bases. We confirmed this finding by sequencing two blood samples originating from human patients which showed different levels of parasitaemia. The estimated enrichment for patient samples (3.9 and 5.8 fold) was in line with the enrichment observed for diluted samples with low percentages of *P. falciparum* DNA (3.3-5.1 fold). Overall, the degree of enrichment for all samples was similar to previous reports where adaptive sampling enriched target DNA 2-up to 6-fold in various sample types^11,33–35^.

Sequencing of patient samples in adaptive sampling mode resulted in enough reads to cover at least 97% of the *P. falciparum* reference genome at a mean depth of 20 or 6, depending on the parasitic load of the sample. This allowed addressing common clinical or research questions, since a high concordance of drug resistance SNP calls obtained via Nanopore and Sanger sequencing was found, showing that a median sequencing depth of 5 is already sufficient to confidently call SNPs with Nanopore (kit SQK-LSK114). It should be noted, however, that the MOI for patient sample 2 with the lowest amount of parasite DNA was 1. A higher MOI may complicate interpretation of the results and would require a higher sequencing depth. In addition, draft assemblies could be constructed, consisting of 138 and 198 contigs. This is sufficient to recover most *Plasmodium* genes, but a higher sequencing depth would be required to generate complete, chromosome-level *de novo* assemblies.

In comparison to existing WGS methods relying on parasite enrichment (sWGA, hybrid selection), the nanopore adaptive sampling method presented here can deliver sequencing results sooner after obtaining the blood sample, and increases sample preparation flexibility. Although leukocyte depletion (cellulose columns or Plasmodipur filter) only adds 30min-3h to the total sample processing time, it requires fresh blood samples^7^. Therefore, this method is only suited when a lab facility is near to the sampling site, and has the capacity to process samples on arrival. sWGA, hybrid selection or leukocyte depletion methods result in a higher parasite enrichment than observed for nanopore adaptive sampling in this study^3–5^, and are more cost-efficient at the moment of writing. However, the MinION sequencing device itself is cheap compared to the cost of purchasing other sequencing platforms or repeatedly shipping and outsourcing to a sequencing facility. In addition, on-site sequencing in endemic areas can increase country-ownership of the data, and autonomy of local research groups.

The Nanopore adaptive sampling workflow is less complicated than other existing methods, compatible with frozen blood, and therefore more suited for low resource settings. To further improve field applicability, up to 500μL patient blood could be collected by finger prick in an EDTA microtainer, instead of by intravenous puncture as done in this study. The ratio of parasite to human DNA could be improved by spinning down the fresh blood sample upon arrival in the lab, to remove plasma and buffy coat, as opposed to the frozen whole blood sample that was used here. DNA extraction protocols could be optimized to obtain longer fragments, or deplete short fragments, since longer DNA fragments can lead to higher enrichment via adaptive sampling^11^. Algorithmic improvements that make the decision to eject a read faster or more accurate, may further improve the adaptive sampling efficiency.

The method presented here has the potential to evolve into a species-agnostic diagnostics tool. We detected *Toxoplasma gondii* DNA in the studied patient samples, illustrating that lower parasitaemia samples than those used in this study can be used for species identification. Genomic co-characterisation of pathogens infecting a patient could improve pathogen surveillance and inform patient treatment. Clinically distinguishing between severe malaria and bacteraemia or other febrile illnesses is difficult, especially in low-resource settings^36-37^. Malaria also occurs as a co-infection with human African trypanosomiasis which requires a different treatment regimen^38^. Further research is required to investigate whether adaptive sampling-enhanced Nanopore sequencing directly from human blood can detect low-abundant DNA in the sample such as DNA originating from bacteria or viruses.

In conclusion, we showed that adaptive-sampling-enhanced Nanopore sequencing is a promising method to whole genome sequence *P. falciparum* directly from blood samples. It shortens the time to retrieve a *P. falciparum* genome from a human blood sample compared to existing methods, can be deployed in most clinical settings including in field hospitals, and generally provides enough sequencing depth to address common research questions, including calling drug resistance loci. Currently, the high cost per sample is still prohibitive for many field applications, and for lower parasitaemia patient samples (<0.1%) depth of coverage might still be a bottleneck for accurate SNP calling. However, over the course of this study, new ONT chemistry already tripled our output of *P. falciparum* bases for the same price. With future improvements in mind, adaptive Nanopore sequencing holds promise to become a field-deployable tool for rapid *Plasmodium* sequencing.

## Declarations

### Data availability

The data for this study have been deposited in the European Nucleotide Archive (ENA) at EMBL-EBI under accession number PRJEB57715 (https://www.ebi.ac.uk/ena/browser/view/PRJEB57715). Fastq files originating from patient samples were purged from human reads for ethical reasons.

## Acknowledgements

We thank the Flemish Institute for Biotechnology (VIB) neuromics core facility for providing the Fragment Analyser service, Pieter Guetens (ITM) for patient sample genotyping by PCR.

## Funding

This work is supported by the University of Antwerp Industrial Research Fund (IOF) SBO project LeapSEQ, the University of Antwerp Core Facility Funding for BIOMINA, and the Research Foundation Flanders (1S48419N scholarship to KDM), and funding of the Malariology Unit (to ARU). The computational resources and services used in this work were provided by the HPC core facility CalcUA of the University of Antwerp, and VSC (Flemish Supercomputer Center), funded by the Research Foundation Flanders (FWO) and the Flemish Government.

### Ethical approval

This study was approved by the ITM Institutional Review Board (project code 1606) and the Ethics Committee of the UZA and the University of Antwerp (project code 3574).

### Authors’ contributions

BC, KL, KDM and WLC conceived the study. ARU provided patient samples, WLC and KDM performed the analysis and wrote the first draft of the manuscript. BC and KL supervised the study and acquired funding for the study. BC, KL and ARU revised the manuscript. All authors read and approved the final manuscript.

