## Supplementary data for "Selective whole-genome sequencing of *Plasmodium* parasites directly from blood samples by Nanopore adaptive sampling"

### Supplementary Figures

A

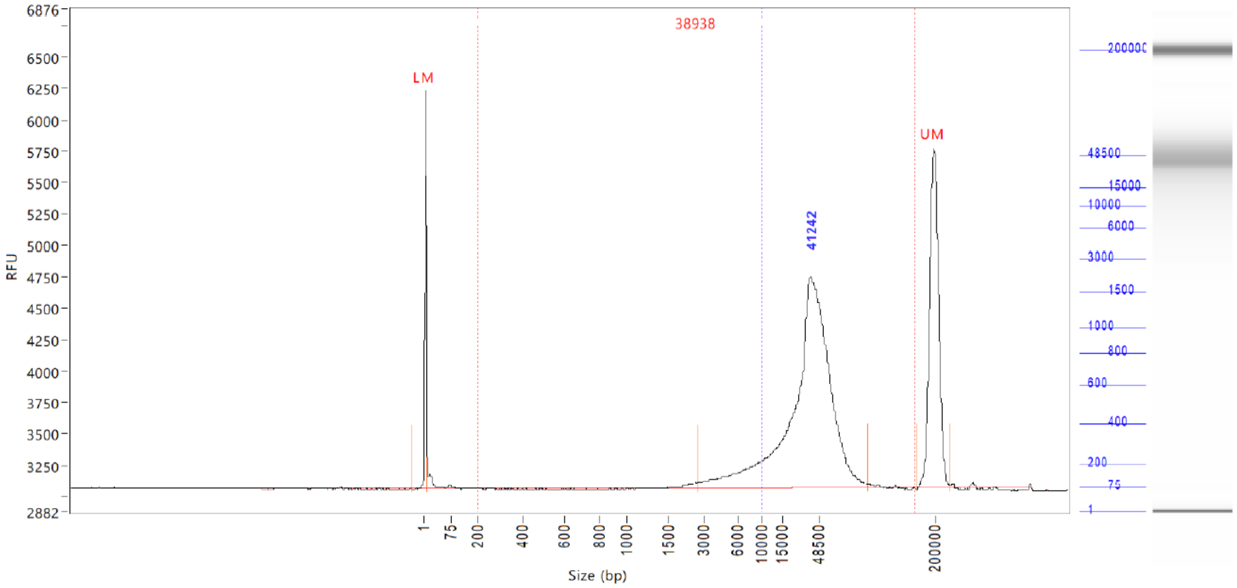


B

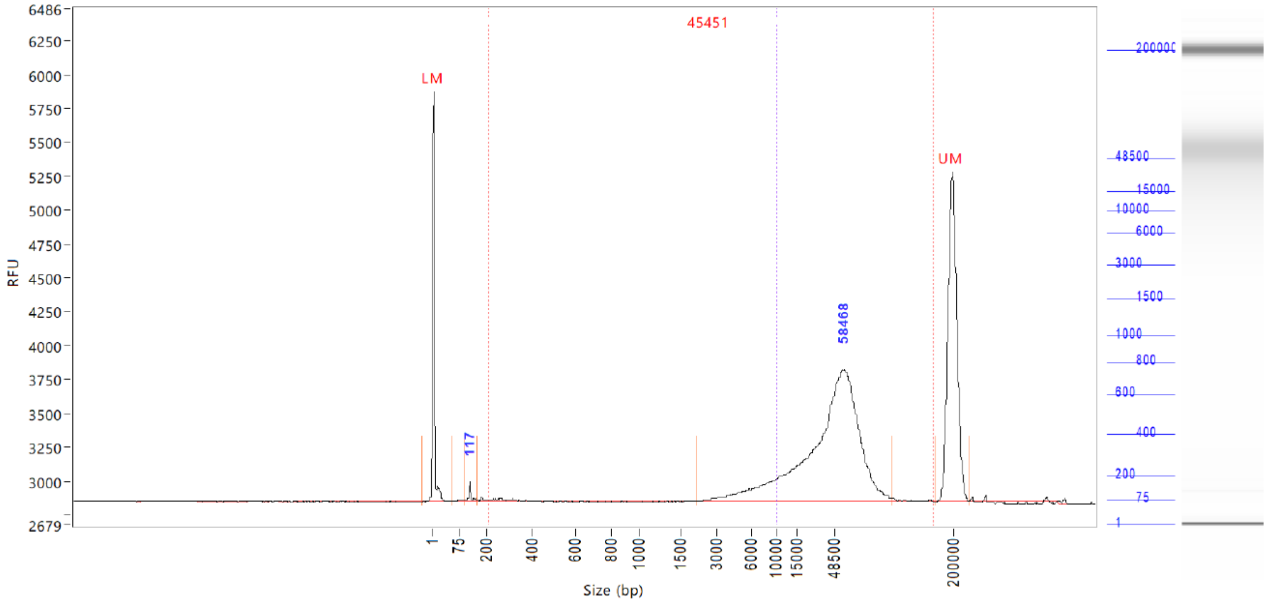


**Supplementary Figure 1. A.** Fragment Analyser plot for commercial human DNA Promega, G304A. **B.** Fragment analyser plot for DNA originating from in vitro cultured *P. falciparum* strain 3D7. DNA was extracted using QIAamp DNA Mini kit (Qiagen) after human white blood cell depletion using Plasmodipur filtration.

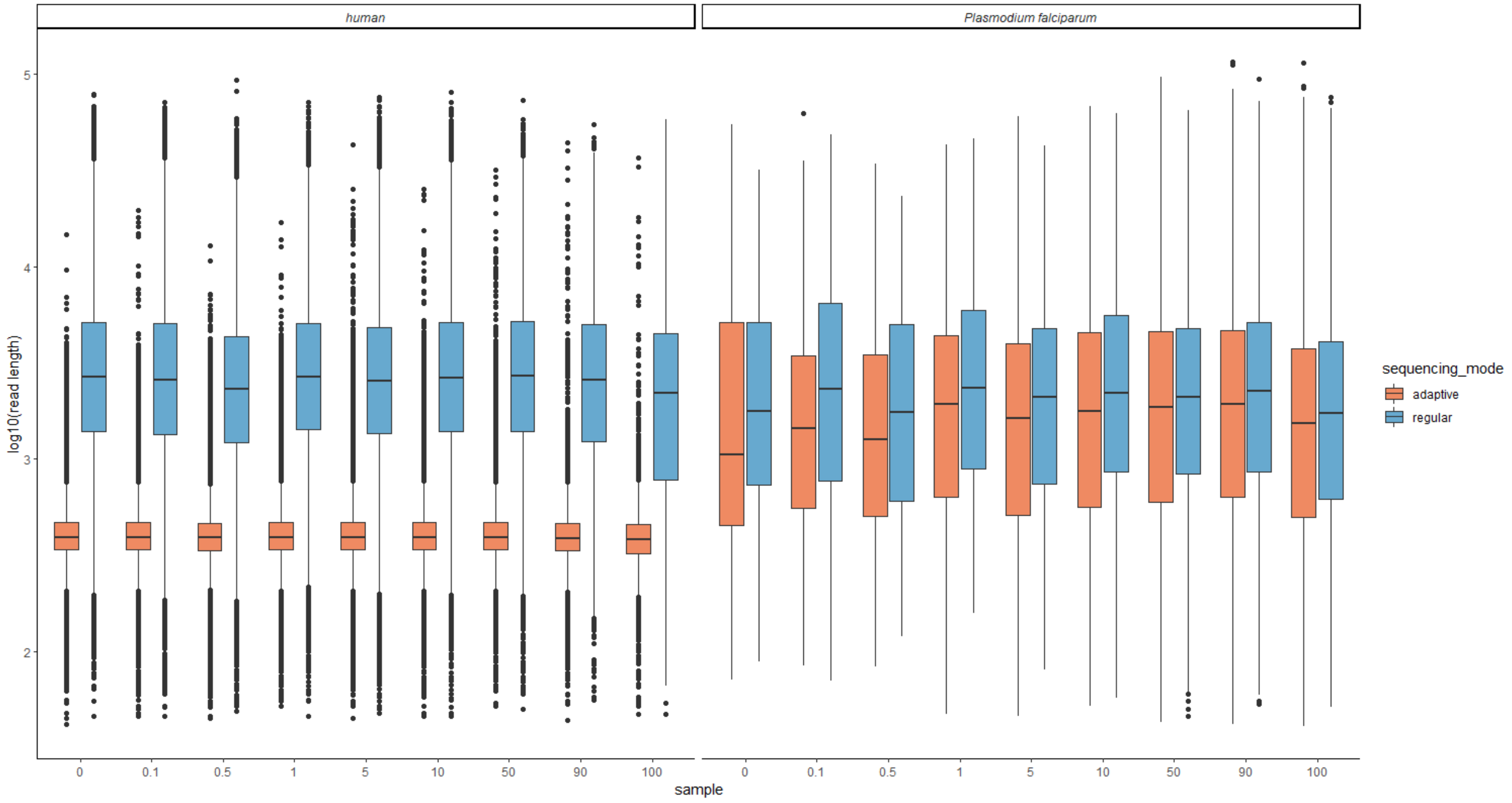


**Supplementary Figure 2.** Short read lengths dominated the read length distributions for human reads in adaptive sampling mode. Log-scaled read length distributions for samples sequenced in either adaptive mode or regular mode (orange and blue respectively) for reads that are either mapped to the human (left-panel) or *P. falciparum* (right panel) genome.

### Supplementary Tables

**Supplementary Table 1.** Comparison of Ligation Sequencing Kit SQK-LSK112 and SQK-LSK114, operating in adaptive mode (enriching for the *P. falciparum* 3D7 genome). Pf = *P. falciparum*.

<table>
<tr><th></th><th></th><th>SQK-LSK112</th><th>SQK-LSK114</th><th>Increase</th></tr>
<tr><td rowspan="3">Patient 1</td><td>mapped bases</td><td>2.6E+09</td><td>7.0E+09</td><td>2.7x</td></tr>
<tr><td>mapped Pf bases</td><td>1.5E+08</td><td>4.7E+08</td><td rowspan="2">3.2x</td></tr>
<tr><td>mean depth Pf genome</td><td>6.3</td><td>20.0</td></tr>
<tr><td rowspan="3">Patient 2</td><td>mapped bases</td><td>3.1E+09</td><td>7.8E+09</td><td>2.6x</td></tr>
<tr><td>mapped Pf bases</td><td>3.3E+07</td><td>1.3E+08</td><td rowspan="2">4.1x</td></tr>
<tr><td>mean depth Pf genome</td><td>1.4</td><td>5.7</td></tr>
</table>

**Supplementary Table 2.** At least three times more reads were sequenced in adaptive sampling mode versus regular sequencing mode.

| **Sample (% *P. falciparum* DNA)** | **Number of reads adaptive sampling** | **Number of reads regular sequencing mode** | **ratio** |
|---|---|---|---|
| 0 | 236576 | 53857 | 4.40 |
| 0.1 | 141183 | 43352 | 3.26 |
| 0.5 | 176857 | 54572 | 3.24 |
| 1 | 149708 | 45870 | 3.26 |
| 5 | 145759 | 43860 | 3.32 |
| 10 | 142062 | 44220 | 3.21 |
| 50 | 92459 | 28297 | 3.27 |
| 90 | 61378 | 19351 | 3.17 |
| 100 | 76389 | 24065 | 3.18 |

**Supplementary Table 3.** Reads mapping to the *P. falciparum* genome with additional blastn hits to the human reference genome. For these reads, the top hits are shown (maximum five, shaded similarly per read ID).

| sequencing mode | read ID | species | percentage identity | E value | bitscore | query length | query coverage |
|---|---|---|---|---|---|---|---|
| regular | 2987f165-274e-4688-a0ff-75353e84b75d | *P. falciparum* | 90.938 | 0 | 5583 | 7713 | 86 |
| regular | 2987f165-274e-4688-a0ff-75353e84b75d | *P. falciparum* | 89.505 | 0 | 3236 | 7713 | 86 |
| regular | 2987f165-274e-4688-a0ff-75353e84b75d | *H. sapiens* | 95.4 | 0 | 918 | 7713 | 7 |
| regular | 43655e5e-1486-4859-806e-ef13a9228c51 | *P. falciparum* | 98.116 | 0 | 3031 | 1770 | 98 |
| regular | 43655e5e-1486-4859-806e-ef13a9228c51 | *P. falciparum* | 98.116 | 0 | 3031 | 1770 | 98 |
| regular | 43655e5e-1486-4859-806e-ef13a9228c51 | *P. falciparum* | 88.374 | 0 | 1539 | 1770 | 73 |
| regular | 43655e5e-1486-4859-806e-ef13a9228c51 | *P. falciparum* | 86.685 | 0 | 1153 | 1770 | 77 |
| regular | 43655e5e-1486-4859-806e-ef13a9228c51 | *P. falciparum* | 87.025 | 3.83E-91 | 342 | 1770 | 77 |
| regular | 5ec63db6-eef3-4aab-b5f0-c3039c4ac906 | *P. falciparum* | 97.323 | 0 | 8148 | 12428 | 75 |

| | | | | | | | |
|---|---|---|---|---|---|---|---|
| regular | 5ec63db6-eef3-4aab-b5f0-c3039c4ac906 | *P. falciparum* | 93.556 | 0 | 5825 | 12428 | 75 |
| regular | 5ec63db6-eef3-4aab-b5f0-c3039c4ac906 | *P. falciparum* | 87.068 | 0 | 817 | 12428 | 75 |
| regular | 5ec63db6-eef3-4aab-b5f0-c3039c4ac906 | *H. sapiens* | 98.235 | 0 | 5528 | 12428 | 25 |
| regular | 5ec63db6-eef3-4aab-b5f0-c3039c4ac906 | *H. sapiens* | 98.11 | 0 | 5507 | 12428 | 25 |
| regular | 7ab3de2f-d63f-413c-b80e-6c359b8c71d6 | *P. falciparum* | 97.722 | 0 | 4711 | 4785 | 57 |
| regular | 7ab3de2f-d63f-413c-b80e-6c359b8c71d6 | *H. sapiens* | 98.519 | 0 | 2846 | 4785 | 43 |
| regular | 7ab3de2f-d63f-413c-b80e-6c359b8c71d6 | *H. sapiens* | 95.503 | 0 | 728 | 4785 | 43 |
| regular | 7ab3de2f-d63f-413c-b80e-6c359b8c71d6 | *H. sapiens* | 80.687 | 5.23E-39 | 171 | 4785 | 43 |
| regular | 7ab3de2f-d63f-413c-b80e-6c359b8c71d6 | *H. sapiens* | 81.69 | 2.43E-37 | 165 | 4785 | 43 |
| regular | ba196188-cdbc-46f5-9437-1d37e7c13cdc | *P. falciparum* | 94.118 | 0 | 26278 | 31976 | 86 |
| regular | ba196188-cdbc-46f5-9437-1d37e7c13cdc | *P. falciparum* | 92.534 | 0 | 5144 | 31976 | 86 |
| regular | ba196188-cdbc-46f5-9437-1d37e7c13cdc | *P. falciparum* | 93.072 | 0 | 4475 | 31976 | 86 |

| | | | | | | | |
|---|---|---|---|---|---|---|---|
| regular | ba196188-cdbc-46f5-9437-1d37e7c13cdc | *P. falciparum* | 92.066 | 0 | 4150 | 31976 | 86 |
| regular | ba196188-cdbc-46f5-9437-1d37e7c13cdc | *P. falciparum* | 94.181 | 0 | 1507 | 31976 | 86 |
| adaptive | 4acd0a5f-bc1f-41e5-bdff-7430fe425d4a | *P. falciparum* | 97.001 | 0 | 1609 | 1709 | 99 |
| adaptive | 4acd0a5f-bc1f-41e5-bdff-7430fe425d4a | *P. falciparum* | 90.621 | 0 | 1003 | 1709 | 99 |
| adaptive | 4acd0a5f-bc1f-41e5-bdff-7430fe425d4a | *H. sapiens* | 100 | 6.98E-04 | 52.8 | 1709 | 2 |
| adaptive | 5c558d0d-c955-474e-aaca-92a4036f04e1 | *P. falciparum* | 97.836 | 0 | 23865 | 27188 | 91 |
| adaptive | 5c558d0d-c955-474e-aaca-92a4036f04e1 | *P. falciparum* | 97.279 | 0 | 19224 | 27188 | 91 |
| adaptive | 5c558d0d-c955-474e-aaca-92a4036f04e1 | *P. falciparum* | 93.122 | 6.16E-70 | 276 | 27188 | 91 |
| adaptive | 5c558d0d-c955-474e-aaca-92a4036f04e1 | *P. falciparum* | 92.157 | 1.36E-51 | 215 | 27188 | 91 |
| adaptive | 5c558d0d-c955-474e-aaca-92a4036f04e1 | *P. falciparum* | 100 | 5.22E-06 | 63.9 | 27188 | 91 |
| adaptive | a4fb204e-26ba-46fe-bc73-f9549cddde4f | *P. falciparum* | 97.561 | 0 | 26195 | 17432 | 99 |
| adaptive | a4fb204e-26ba-46fe-bc73-f9549cddde4f | *P. falciparum* | 86.202 | 0 | 2278 | 17432 | 99 |

| | | | | | | | |
|---|---|---|---|---|---|---|---|
| adaptive | a4fb204e-26ba-46fe-bc73-f9549cddde4f | *H. sapiens* | 100 | 0.007 | 52.8 | 17432 | 0 |
| adaptive | b8297f1a-711a-447e-b282-36d4f802b6a6 | *P. falciparum* | 100 | 0 | 1210 | 655 | 100 |
| adaptive | b8297f1a-711a-447e-b282-36d4f802b6a6 | *P. falciparum* | 100 | 0 | 1210 | 655 | 100 |
| adaptive | b8297f1a-711a-447e-b282-36d4f802b6a6 | *P. falciparum* | 99.39 | 0 | 1186 | 655 | 100 |
| adaptive | b8297f1a-711a-447e-b282-36d4f802b6a6 | *P. falciparum* | 92.519 | 0 | 918 | 655 | 100 |
| adaptive | b8297f1a-711a-447e-b282-36d4f802b6a6 | *P. falciparum* | 92.519 | 0 | 918 | 655 | 100 |
| adaptive | d59829de-a98a-4b72-9cea-4fff17168c3c | *P. falciparum* | 96.114 | 0 | 1075 | 1262 | 96 |
| adaptive | d59829de-a98a-4b72-9cea-4fff17168c3c | *P. falciparum* | 85.235 | 5.32E-158 | 564 | 1262 | 96 |
| adaptive | d59829de-a98a-4b72-9cea-4fff17168c3c | *H. sapiens* | 100 | 5.12E-04 | 52.8 | 1262 | 2 |

**Supplementary Table 4.** Taxonomic read classification of reads from patient sample 1 and patient sample 2. Reads were compared to the PlusPF-16 database using kraken2. Species for which both samples contained less than 10 reads were not listed.

| **taxon ID** | **Taxon name** | **Patient sample 1 count** | **Patient sample 2 count** |
|---|---|---|---|
| 9606 | *Homo sapiens* | 6727046 | 11328372 |
| 36329 | *Plasmodium falciparum* 3D7 | 59407 | 18594 |
| 508771 | *Toxoplasma gondii* ME49 | 10487 | 22467 |
| 2759 | Eukaryota | 2229 | 3870 |
| 5854 | *Plasmodium reichenowi* | 753 | 103 |
| 418107 | *Plasmodium* (*Laverania*) | 684 | 93 |
| 131567 | cellular organisms | 642 | 1003 |
| 208452 | *Plasmodium coatneyi* | 195 | 68 |
| 54757 | *Plasmodium vinckei vinckei* | 194 | 72 |
| 647221 | *Plasmodium gaboni* | 117 | 11 |
| 5823 | *Plasmodium berghei* ANKA | 112 | 35 |
| 5855 | *Plasmodium vivax* | 101 | 27 |
| 880535 | *Plasmodium* sp. *gorilla* clade G2 | 91 | 21 |
| 418101 | *Plasmodium* (*Vinckeia*) | 50 | 11 |
| 31271 | *Plasmodium chabaudi chabaudi* | 39 | 10 |
| 5820 | *Plasmodium* | 27 | <10 |
| 5861 | *Plasmodium yoelii* | 21 | <10 |
| 418103 | *Plasmodium* (*Plasmodium*) | 18 | <10 |
| 13502 | *Brettanomyces nanus* | 16 | 22 |
| 5851 | *Plasmodium knowlesi* strain H | 15 | <10 |
| 85471 | *Plasmodium relictum* | 15 | <10 |
| 451864 | Dikarya | 13 | 18 |
| 1747 | *Cutibacterium acnes* | 12 | 23 |
| 280036 | *Sporisorium graminicola* | 10 | 24 |
| 367775 | *Cryptococcus gattii* WM276 | <10 | 10 |

| | | | |
|---|---|---|---|
| 352472 | *Dictyostelium discoideum* AX4 | <10 | 13 |

**Supplementary Table 5.** Mean and median depth for reads from patient sample 1 mapped to the *P. falciparum* 3D7 reference genome, and the percentage of the 3D7 genome that is covered.

| | mean depth | median depth | coverage (%) |
|---|---|---|---|
| **full genome** | 19.7 | 20 | 99.8 |
| chromosome 1 | 19.6 | 19 | 99.5 |
| chromosome 2 | 19.9 | 20 | 99.8 |
| chromosome 3 | 20.2 | 20 | 99.9 |
| chromosome 4 | 19.1 | 19 | 99.5 |
| chromosome 5 | 19.2 | 19 | 99.9 |
| chromosome 6 | 19.7 | 20 | 99.6 |
| chromosome 7 | 18.4 | 19 | 99.1 |
| chromosome 8 | 19.9 | 20 | 99.9 |
| chromosome 9 | 19.6 | 19 | 99.9 |
| chromosome 10 | 19.4 | 19 | 99.9 |
| chromosome 11 | 20.2 | 20 | 99.8 |
| chromosome 12 | 19.0 | 19 | 99.8 |
| chromosome 13 | 20.3 | 20 | 99.9 |
| chromosome 14 | 19.6 | 20 | 99.9 |
| apicoplast | 14.8 | 15 | 99.9 |
| mitochondrial plasmid | 520.8 | 522 | 100.0 |

**Supplementary Table 6.** Mean and median depth for reads from patient sample 2 mapped to the *P. falciparum* 3D7 reference genome, and the percentage of the 3D7 genome that is covered.

| | **mean depth** | **median depth** | **coverage (%)** |
|---|---|---|---|
| **full genome** | 5.6 | 5 | 97.4 |
| chromosome 1 | 6.1 | 6 | 97.0 |
| chromosome 2 | 5.2 | 5 | 97.2 |
| chromosome 3 | 6.0 | 5 | 97.4 |
| chromosome 4 | 5.3 | 5 | 94.7 |
| chromosome 5 | 5.2 | 5 | 98.2 |
| chromosome 6 | 5.5 | 5 | 96.9 |
| chromosome 7 | 5.6 | 5 | 95 |
| chromosome 8 | 5.7 | 6 | 95.7 |
| chromosome 9 | 5.5 | 5 | 98.1 |
| chromosome 10 | 5.4 | 5 | 97.8 |
| chromosome 11 | 5.5 | 5 | 97.9 |
| chromosome 12 | 5.9 | 6 | 96.8 |
| chromosome 13 | 5.5 | 5 | 98.6 |
| chromosome 14 | 5.7 | 6 | 98.4 |
| apicoplast | 6 | 6 | 100 |
| mitochondrial plasmid | 123.5 | 125 | 100.0 |

**Supplementary Table 7.** The number of contigs assigned to each chromosome for Flye genome assemblies of patient 1 and 2.

| | Number of assigned contigs | |
|---|---|---|
| | **Patient 1** | **Patient 2** |
| chromosome 1 | 2 | 10 |
| chromosome 2 | 5 | 10 |
| chromosome 3 | 1 | 6 |
| chromosome 4 | 9 | 11 |
| chromosome 5 | 1 | 10 |
| chromosome 6 | 4 | 6 |
| chromosome 7 | 6 | 11 |
| chromosome 8 | 7 | 9 |
| chromosome 9 | 6 | 17 |
| chromosome 10 | 4 | 11 |
| chromosome 11 | 5 | 20 |
| chromosome 12 | 8 | 15 |
| chromosome 13 | 6 | 26 |
| chromosome 14 | 3 | 24 |
| apicoplast | 1 | 1 |
| mitochondrial plasmid | 1 | 1 |
| unassigned | 69 | 10 |

**Supplementary Table 8 (provided as separate dataset).** Total sequencing depth and depth of the resistance allele for drug-resistance associated markers in *P. falciparum*[1] in patient 1 and 2. The higher multiplicity of infection (MOI) of patient 1 explains the presence of both resistant and non-resistant alleles at some loci. For 38 of the resistance associated markers, a PCR fragment was Sanger sequenced to assess whether the SNP was called correctly with nanopore sequencing.
